# Sex-Specific Cross Tissue Meta-Analysis Identifies Immune Dysregulation in Women with Alzheimer’s Disease

**DOI:** 10.1101/2020.04.24.060558

**Authors:** Manish D Paranjpe, Stella Belonwu, Jason K Wang, Tomiko Oskotsky, Aarzu Gupta, Alice Taubes, Kelly Zalokusky, Ishan Paranjpe, Benjamin S Glicksberg, Yadong Huang, Marina Sirota, for the Alzheimer’s Disease Neuroimaging Initiative

## Abstract

In spite of evidence of females having a greater lifetime risk of developing Alzheimer’s Disease (AD) and greater apolipoprotein E4-related (apoE4) AD risk compared to males, molecular signatures underlying these findings remain elusive. We took a meta-analysis approach to study gene expression in the brains of 1,084 AD patients and age-matched controls and whole blood from 645 AD patients and age-matched controls. Gene-expression, network-based analysis and cell type deconvolution approaches revealed a consistent immune signature in the brain and blood of female AD patients that was absent in males. Machine learning-based classification of AD using gene expression from whole blood in addition to clinical features revealed an improvement in classification accuracy upon stratifying by sex, achieving an AUROC of 0.91 for females and 0.80 for males. These results help identify sex and apoE4 genotype-specific transcriptomic signatures of AD and underscore the importance of considering sex in the development of biomarkers and therapeutic strategies for AD.

## INTRODUCTION

Alzheimer’s disease (AD) is a progressive neurodegenerative disorder and the most common cause of dementia^1, 2^. It is pathologically characterized by the deposition of extracellular amyloid β (Aβ) and intracellular tau, otherwise referred to as plaques and neurofibrillary tangles, respectively^3–5^. AD is also marked by neuronal loss, impaired neurotransmitter signaling, neuroinflammation, and dysregulation of neuronal metabolism and immune response in the central nervous system^6–8^. AD prevalence increases dramatically with age, where the majority of cases are in individuals above the age of 65^1, 9^. Although AD was identified more than a century ago^10^, its cause and pathophysiology are not fully understood, and there are no available treatments that aid in halting or reversing the disease^11^. Accordingly, it is of high priority to tackle AD, as it is projected to triple in incidence by 2050 as a consequence of population aging^6, 8, 12^ and, to date, has no disease-modifying therapies.

While the exact cause and pathophysiology remain unknown, a number of mutations and genetic risk factors have been identified as associated with AD. Apolipoprotein E (apoE) is the most common genetic risk factor for late onset AD^8, 13–18^. ApoE is a lipid binding protein, that plays a central role in lipid transport and metabolism. It is highly expressed in the brain, and is important for maintaining neuronal membranes during inflammation and damage. In humans, apoE has three isoforms, apoE2, apoE3, and apoE4, which are encoded by the three alleles, ε2, ε3, and ε4, of the apoE gene, respectively. The ε2 isoform has been shown to be protective against AD, while the ε4 isoform (apoE4) is associated with increasing the risk and lowering the age of onset for developing late onset AD in a gene dose-dependent manner^19, 20^. Specifically, one copy of the ε4 isoform confers a 3 to 4-fold increased risk and 7 year decrease in age of onset, while two copies confers a 12 to 15-fold increased risk of AD, and a 14 year decrease in age of onset^8, 21^.

Sex is another major risk factor in AD. Female sex is associated with increased AD incidence, exacerbated pathophysiology and increased rate of cognitive decline related to the disease progression^8, 22–25^. It has been conjectured that the higher prevalence in females is a result of longer life span^8, 25^. Alternatively, studies have alluded to sex-specific hormonal and metabolic changes that interplay with the onset and progression of AD dementia^6, 16, 26^. Sex also interacts with apoE isoform status, where females with the apoE4 isoform are at increased risk compared to males^27–29^. Despite the clear therapeutic potential to better understanding these pathophysiological patterns, there is still little understanding of the mechanisms underlying sex-specific differences in AD.

With the rising prevalence of AD, it is critical to facilitate the development of robust means to detect AD early and discover therapeutic interventions^30–33^. Technological innovations and the increasing availability of large transcriptomic datasets present worthwhile avenues to study and characterize the molecular underpinnings of AD stratified by sex. Here, we analyze publicly available gene expression datasets from over 1,500 brain and blood samples to characterize this highly complex disease. To derive sex-specific transcriptomic molecular signatures, we perform a meta-analysis, differential gene expression, weighted gene co-expression network analysis, pathway enrichment, and cell-type deconvolution in a large cohort of brain and blood samples from AD patients and healthy controls (Figure 1). We further characterize these signatures and apply machine learning to build a predictive model based on biomarkers identified in the blood of AD patients. Our findings reveal underlying mechanisms of sex differences, which provide clinical implications for identifying more accurate, and less invasive biomarkers, as well as efficacious therapeutics tailored to better fit the complex molecular profiles in AD.

**Figure 1:**
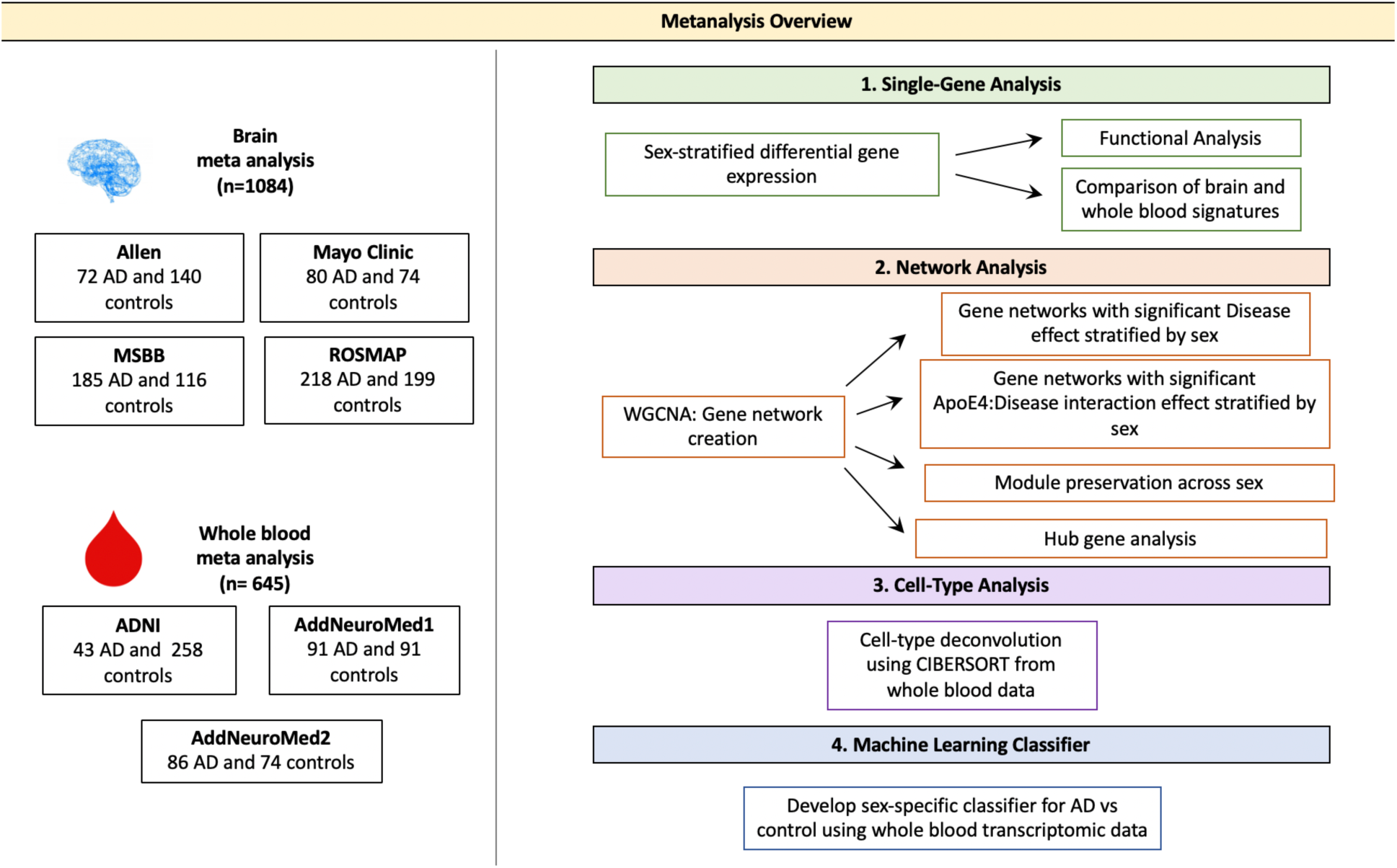
Meta-analysis Overview. Diagram depicting the study overview including all datasets used and analyses performed. Data sets were obtained via searching GEO or PubMed for the keyword Alzheimer’s Disease. Samples with neurological conditions other Alzheimer’s, including Parkinson’s Disease and Huntington’s Disease, and single cell preparations were excluded from analysis. Datasets were merged using the ComBat package in R. WGCNA was used for network analyses. CIBERSORT was used for cell type deconvolution. The linear SVM was trained to classify AD and control patients using the transcriptomic signature obtained via meta-analysis of blood studies. The performance of a molecular model consisting of gene expression, age, sex and apoE4 status was compared to that clinical model with age, sex and apoE4 status as features.

## METHODS

### Study Cohorts

Publicly available RNA-sequencing (RNA-Seq) and microarray datasets from the Gene Expression Omnibus (GEO) and from consortium studies indexed on PubMed were searched for the key word “Alzheimer’s”. To minimize technical variability, brain samples were restricted to RNA-sequencing studies while blood analyses were restricted to microarray studies. Samples were curated to include bulk gene expression from subjects with Alzheimer’s or elderly healthy individuals with no history of neurodegenerative disease. Individuals with non-Alzheimer’s neurodegenerative diseases including Huntington’s and Parkinson’s were excluded. Brain samples were restricted to the hippocampus, parietal cortex, temporal cortex and prefrontal cortex. Additional clinical covariates, including age, sex, apoE4 carrier status, education were recorded for the samples and used as covariates or stratification variables in subsequent analyses.

### Gene Expression Meta-Analysis

Meta-analysis was conducted separately for brain and blood studies according to standard quality control, normalization, and batch correction procedures. All data processing was conducted using R (v3.6.1).

#### Brain studies

Raw RNA-sequencing data were processed for the Mount Sinai Brain Bank (MSBB)^34^, Mayo Clinic RNAseq ^35^, and Religious Orders Study and Memory and Aging Project (ROSMAP)^36^ as previously described in the AMP-AD consortium project. Briefly, read alignment and counting was performed using STAR^38^. Alignment quality metrics were generated using PICARD^39^. For the Allen dataset, expected counts produced using RSEM were downloaded from the Allen Brain Atlas: Aging Dementia and TBI Study website^40^. Counts-per-million (CPM) were calculated for all studies. Genes with less than 1 CPM in at least 50% of samples across tissue diagnosis group were removed. Genes with missing gene length or GC content percentage metrics were removed. Library normalization was performed using conditional quantile normalization.

Following read alignment and normalization, studies were merged using common genes between the four studies. Mean value imputation was performed for missing gene expression values. Quantile normalization was performed across studies. The ComBat function from the *sva* package^41^ was used to perform cross-study normalization, retaining variation in apoE4 carrier status, sex, and diagnosis. Principal component analysis (PCA) plots were generated to evaluate successful batch correction and to detect outliers.

#### Blood studies

Study data were downloaded from GEO for the AddNeuroMed datasets^42^ or the Alzheimer’s Disease Neuroimaging Initiative Consortium^43^ (ADNI) for the ADNI dataset and processed. Raw data were not available for the ADNI dataset and therefore normalized expression data were used for all studies. Outlier removal was performed on individual studies by removing probes whose mean expression was outside 1.5 times the interquartile range. Probe IDs were mapped to gene symbols. Expression value of probes mapping to the same gene were reported as the median of all probes mapping to that gene^44^. Quantile normalization was performed across studies. Similar to the brain data analysis, the ComBat function from the *sva* package was used to perform cross-study normalization, retaining variation in apoE4 carrier status, sex and diagnosis. Principal component analysis (PCA) plots were generated to evaluate successful batch correction.

### Differential Gene Expression Analysis

All differential gene expression analyses were performed separately for brain and blood samples. The Limma package^45^ was used to determine differentially expressed genes between cases and controls all together and stratified by sex. In each model, age and apoE4 carrier status were included as covariates to minimize confounding. An additional covariate of education was used in the blood analyses. Education was not available for all brain samples and therefore was not included as a covariate. A cutoff false discovery rate (FDR) of 0.05 and fold change (FC) of greater than or equal to 1.2 was used for brain analyses. Fold changes were calculated using the individual study data before merging and weighted by sample size. For blood analyses, a FC cutoff was not used to maximize gene discovery. Significant overlap between up- and down-regulated genes between males and females was assessed using a hypergeometric test. Functional enrichment analysis of gene lists was carried out by overrepresentation analysis using the KEGG^46^ database of biological pathways.

### Network Analysis

#### Weighted Gene Co-Expression Network Analysis

In order to detect gene network level differences, network analysis was performed using Weighted Gene Co-Expression Network Analysis (WGCNA)^47^. All analyses were performed separately for brain and blood samples. In signed WGCNA, a module is defined as a set of genes whose expression is highly correlated in the same direction. Signed gene co-expression networks were created separately for male and female samples to identify sex-specific gene modules. Module Z-summary scores were computed to assess module preservation between male and female networks, as described previously^48^. A Z-summary score greater than ten was considered to be strong evidence of preservation between the two networks. A score between two and ten was considered to represent weak to moderate evidence of preservation, as previously described^48^.

Association between module gene expression and case/control status was assessed by relating the module eigengenes, defined as the first principal component of the genes in a given module, to case/control status using linear regression. Age, apoE4 carrier status, and education (for blood samples) were used as covariates to minimize confounding. An additional analysis identifying apoE-by-disease interaction effects was performed by adding the interaction term: apoE4 carrier status:case/control status to the previous model. Significant modules were characterized by performing functional gene enrichment using the KEGG database of biological pathways^31^.

#### Hub Gene Analysis

To identify central regulators of gene expression, we identified hub genes within significant modules, as described previously^47^. Hub genes were defined as genes with gene significance (the correlation between the gene expression and case/control status) greater than 0.2 and module membership (the correlation between gene expression and module eigengene) greater than 0.8, as previously described^47^. We also restricted hub genes to those that were differentially expressed in AD vs control. Network visualization using the STRING v11^51^ database was used to assess evidence for protein-protein interactions between hub genes.

### Cell-type Deconvolution

CIBERSORT^52^ was applied to the transcriptomic signatures generated in the blood meta-analysis to deconvolve gene expression data into cell type composition and identify sex-specific dysregulation of immune cell types between cases and controls. CIBERSORT applies a linear support vector regression method to solve the problem: m= f x B where m is an input mixture of gene expression data for a given sample, f is a vector consisting of fractions of each cell type in the mixture and B is a matrix of reference gene expression profiles. A gene expression profile of 22 reference cell populations was built using differential gene expression of purified or enriched cell populations from the authors of CIBERSORT.

CIBERSORT was used to deconvolve gene expression data from pooled male and female data, male only samples, and female only samples. In each condition, differences in cell type proportions between cases and controls were compared using a linear regression model adjusting for age, sex (in the pooled male female analysis), and apoE4 carrier status. An additional analysis identifying apoE4-by-disease interaction effects was performed by adding the interaction term: apoE4 carrier status:case/control status to the previous model. A cutoff FDR of 0.05 was deemed significant.

### Classification of Healthy and Alzheimer’s Disease Patients

A linear support vector machine (SVM) model with *l*_1_ regularization to enforce feature sparsity was used to classify Alzheimer’s patients and healthy controls based on blood gene expression data. To assess the relative value of stratifying by sex in increasing model performance, we compared the performance of three models built using pooled male and female samples, male samples only, and female samples only. We also compared the performance of a ‘clinical model’ with age, sex (for male and female pooled samples), and apoE4 carrier status information to a ‘clinical + molecular model’ which included age, sex (for male and female pooled samples), apoE4 carrier status, and transcriptomic data from the blood meta-analysis.

For each model, data were split into 75% training/validation and 25% test sets using a class balancing procedure to maintain a constant case/control ratio across training/validation and test sets. A random search over the space 10^-4^ to 10^4^ with five-fold cross validation was used to optimize the C hyper-parameter, or the degree of regularization penalty applied for misclassified points. Receiver operating characteristic (ROC) curves were generated from the test set. Model performance was assessed using the area under the ROC curves. Feature importance was determined using the absolute value of the model coefficients corresponding to the vector coordinates orthogonal to the model hyperplane.

## RESULTS

### Study Cohort Characteristics

We obtained four publicly available RNA-seq data sets (Allen Brain Institute Aging Dementia and TBI study, Mayo Clinic RNA-seq, MSBB, and ROSMAP) from the brain (temporal cortex, parietal cortex, prefrontal cortex, and hippocampus) and three microarray datasets from whole blood (AddNeuroMed cohort 1, AddNeuroMed cohort 2 and ADNI). After outlier removal, we included a total of 1,084 brain samples (58% female; 26% apoE4 carriers) and 645 blood samples (58% female; 38% apoE4 carriers) in our analysis. Table 1 shows a summary of sample annotations including number of cases and controls, apoE carrier status, and number of males and females for brain datasets and blood datasets.

**Table 1:** Meta-analysis Study Characteristics

| Study | Accession | Total participants | AD, no. (%) | AD |  | CN |  |
| --- | --- | --- | --- | --- | --- | --- | --- |
|  |  |  |  | Female /Male (% Female) | apoE4 /No (% Yes) | YesFemale /Male (% Female) | apoE4 Yes /No (% Yes) |
| Brain Transcriptomic Studies |  |  |  |  |  |  |  |
| Allen | https://aging.brain-map.org/ | 212 | 72 (34) | 29/43 (40) | 22/50 (31) | 54/86 (39) | 19/121 (14) |
| Mayo Clinic RNA-Seq | syn5550404 | 154 | 80 (52) | 49/31 (61) | 42/38 (53) | 36/38 (49) | 9/65 (12) |
| MSBB | GSE52564 | 301 | 185 (62) | 131/54 (71) | 63/122 (34) | 57/59 (49) | 16/100 (13) |
| ROSMAP | syn3219045 | 417 | 218 (52) | 151/67 (70) | 83/135 (38) | 122/77 (61) | 33/166 (17) |
| Sum |  | 1084 | 555 (52) | 360/195 (65) | 210/345 (38) | 269/260 (51) | 77/452 (15) |
| Whole Blood Transcriptomic Studies |  |  |  |  |  |  |  |
| ADNI | http://adni.loni.usc.edu/ | 301 | 43 (14) | 17/26 (40) | 32/11 (74) | 135/125 (52) | 71/189 (27) |
| AddNeuroMed1 | GSE63060 | 182 | 91 (50) | 65/26 (71) | 52/39 (57) | 55/36 (60) | 30/61 (33) |
| AddNeuroMed2 | GSE63061 | 160 | 86 (43) | 59/27 (69) | 47/39 (55) | 45/29 (61) | 15/59 (20) |
| Sum |  | 645 | 220 (34) | 141/79 (64) | 131/89 (60) | 235/190 (55) | 116/309 (27) |

In the brain datasets, compared to controls, AD patients were significantly older (mean ± SD for AD: 86.5 ± 6.0 years and controls: 84.8 ± 7.4 years; two sample t-test, *P* < 0.001), more likely to be apoE4 carriers (AD: 38% carriers vs controls: 15% carriers; Chi-squared test, *P* < 0.001), and more likely to be females (AD: 65% female vs controls: 51% female; Chi-squared test, *P* < 0.001).

In the blood datasets, compared to controls, AD patients were significantly older (mean ± SD for AD: 77.0 ± 7.1 years and controls: 74.7 ± 5.7 years; two sample t-test, *P* < 0.001), more likely to be apoE4 carriers (AD: 60% carriers vs controls: 27% carriers; Chi-squared test, *P* < 0.001), more likely to be females (AD: 64% female vs controls: 55% female; Chi-squared test, *P* < 0.001), and had more years of education (mean ± SD for AD: 9.4 ± 4.8 years and controls: 13.9 ± 4.7 years; two sample t-test, *P* < 0.001).

Studies were merged and batch corrected using ComBat resulting in 13,500 common genes across 1,084 samples for brain studies and 3,371 common genes across 645 samples for blood studies. Supplementary Figure S1 and S2 show PCA plots before and after batch correction, demonstrating successful data merging and batch effect removal.

### Differential Gene Expression in the Brain Identifies a Distinct Sex-Specific Signature of AD

We observed distinct AD-associated transcriptomic signatures in the brain in males and females. A total of 981 genes were differentially expressed in females, including 583 upregulated genes and 398 downregulated genes (FC > 1.2, q < 0.05; Figures 2A-B; Supplementary Table 1). In males, 513 genes were differentially expressed, including 415 upregulated genes and 98 downregulated genes (FC > 1.2, q < 0.05; Figures 2A-B; Supplementary Table 1). Altogether, 631 genes were uniquely dysregulated in females, including 309 upregulated genes and 327 downregulated genes. In males, 166 genes were uniquely dysregulated, including 141 upregulated genes and 27 downregulated genes. There was a significant overlap of dysregulated genes across males and females (*P* < 0.05; hypergeometric test).

**Figure 2:**
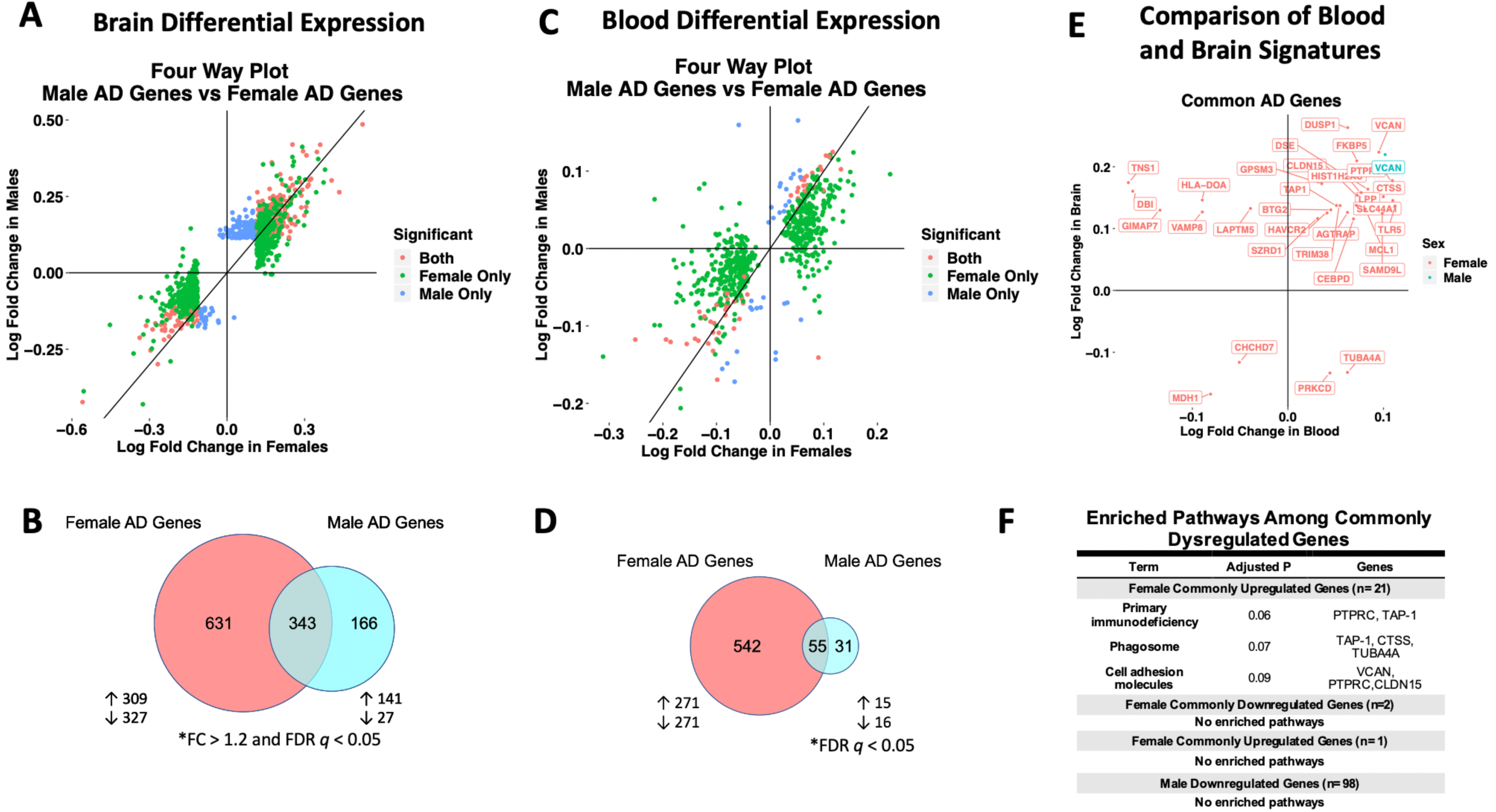
Cross-Tissue Sex Specific Differential Gene Expression. **A.** Four-way plot with fold change in males vs fold change in females depicting differentially expressed genes in the brain. Differential expression was defined using a fold change > 1.2 and FDR *P* < 0.05. Covariates of age and sex were included in statistical analyses. **B.** In the brain, a total of 631 genes were uniquely dysregulated in females with AD while 166 genes were uniquely dysregulated in males with AD. Common to both males and females in the brain were 343 genes. **C.** Four-way plot with fold change in males vs fold change in females depicting differentially expressed genes in the blood. Differential expression was defined using a fold change > 1.2. Covariates of age, sex, and education were included in statistical analyses **D.** A total of 542 genes were uniquely dysregulated in females with AD while 31 genes were uniquely dysregulated in males with AD in blood. Common to both males and females in the brain were 55 genes. **E** Fold change plot depicting genes that are dysregulated in both blood and brain tissues. Genes are colored by sex indicating if the gene is dysregulated in male samples (1 gene; red) or female samples (31 genes; blue). **F.** Enrichment analysis of the commonly dysregulated genes depicted in **E**. An adjusted *P*-value cutoff of 0.1 was used for significance to increase power.

Next, we characterized the transcriptomic signatures observed in the brains of male and female AD patients. In females, among upregulated AD genes, we found 69 enriched pathways, many of them relating to components of the innate and adaptive immune system (Table 2; Supplementary Table 2). Several upregulated HLA system genes including HLA-DPB1, HPA-DRA, HLA-DOA, HLA-DRB5, HLA-DMA, HLA-DPA1 contributed to enrichment of a number of pathways relating to response to infection (Table 3). Components of the complement system including C1QA, C4B, and C4A were also uniquely dysregulated in females (Table 4; Supplementary Table 2). We also observed an enrichment of genes in the MAPK signaling pathway including MRAS, MK2, and MK3. Downregulated AD genes in females were enriched for a number of neurological signaling pathways including synaptic vesicle exocytosis, neuroactive ligand-receptor activation, and GnRH signaling (Table 2; Supplementary Table 3).

**Table 2:**
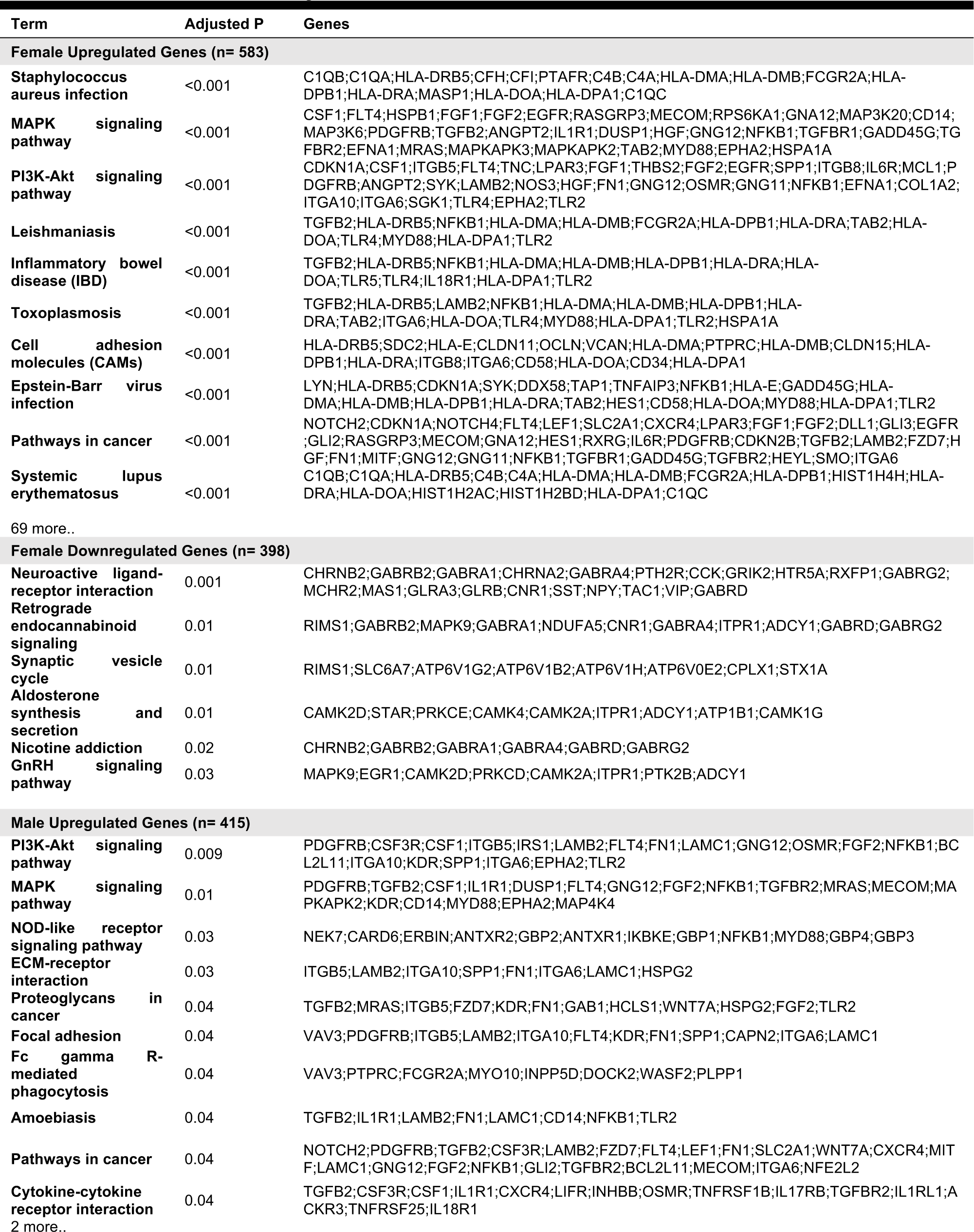
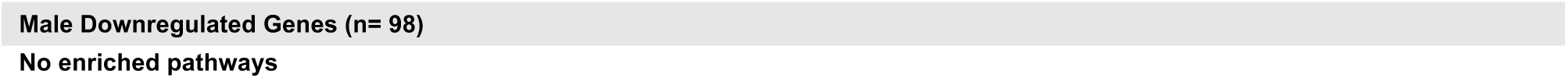
Enriched Pathways in Brain

**Table 3:**
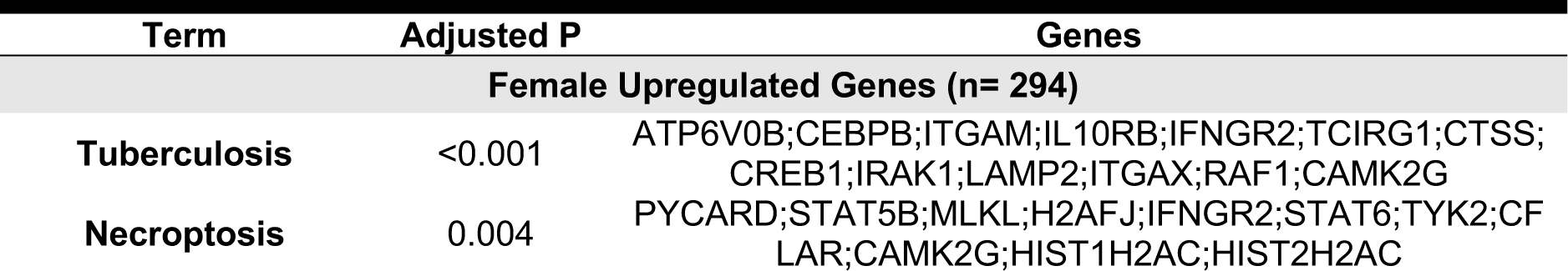
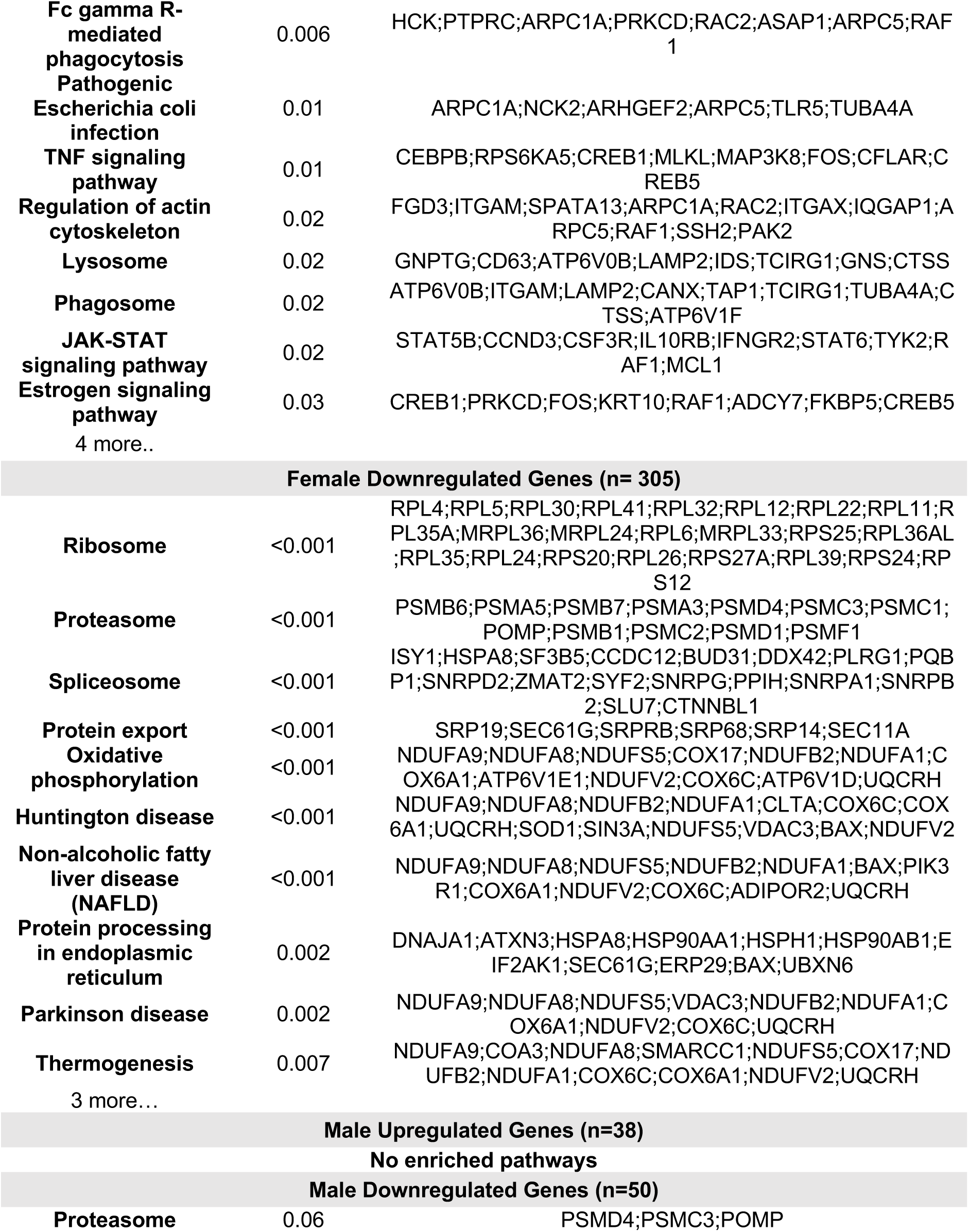
Enriched Pathways in Blood

Strikingly, we observed an enrichment of fewer immune-related pathways in males with AD. Among upregulated genes in male AD patients, we found 12 enriched pathways, including amoebiasis and cytokine-cytokine receptor interaction, suggestive of adaptive and innate immune activation (Table 2; Supplementary Table 4). Similar to females, we also observed an enrichment of the MAPK signaling pathway, including MAP4K4 and MK2, in males. Among downregulated genes in male AD patients, we did not identify significantly enriched pathways. For a full list of enriched pathways, refer to Supplementary Tables S2-S4.

Lastly, we performed a non-stratified analysis comparing gene expression between AD and control samples irrespective of sex. Statistical models were adjusted for sex, apoE4 carrier status, and age. A total of 662 genes were upregulated and 430 genes were downregulated in patients with AD compared to controls (Figure S3, Table 2; Supplementary Table 1. Upregulated genes were enriched for several pathways previously implicated in AD including PI3K-Akt signaling and MAPK signaling as well as a number of immune related pathways including Staphylococcus aureus infection, human papillomavirus infection, and malaria (Supplementary Table S5). Several components of the complement system, including C4B, C4A, C1R, C3AR1, and C5AR1 also contributed to this enrichment (Supplementary Table S6). In our analysis of downregulated genes, we found several pathways related to neuroreceptor signaling and GABAergic transmission were enriched including the genes GABRA1, GNG3, GNG2, SLC32A1, GABRD, and GABRG2 (Supplementary Table S6).

### Network Analysis in the Brain Identifies a Stronger Disease Signature in Females

To assess transcriptomic changes on a gene network level, we utilized WGCNA. Gene networks were derived separately for male and female samples and compared using network preservation methods, as previously described^48^. We identified two AD-associated modules in males and 11 AD-associated modules in females (Figure 3A) that met the significance threshold (FDR < 0.05) and were either positively or negatively correlated with case/control status. Among the male modules, a 463-gene module (termed black) was upregulated in AD, and a 151-gene module (termed tan) was downregulated in AD. The black module in males had significant overlap with two modules in females (termed yellow and pink) (*P* < 0.001; hypergeometric test) as indicated by asterisks in Figure 3B. The black module also had strong preservation in the female network (Z-summary score > 10). Among the female-specific disease associated modules, four modules (termed green, red, black and turquoise) were downregulated in AD, while seven were upregulated (Figure 3A).

**Figure 3:**
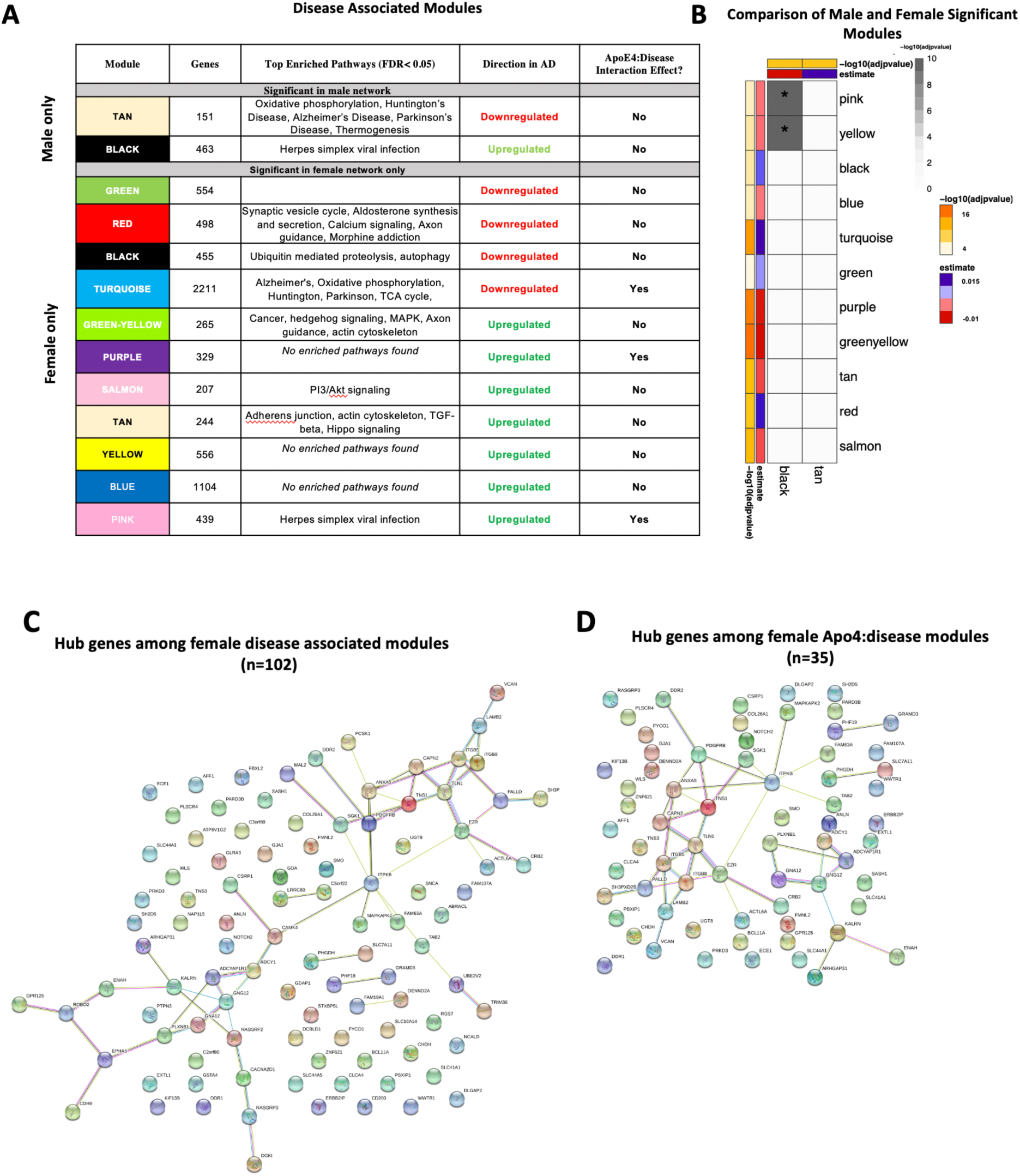
Network Analysis in the Brain. WGCNA was used to construct gene network separately for males and females in the brain. Networks were randomly assigned colors. **A.** A description of the disease-associated gene networks (termed modules) produced using WGCNA. Significant disease-associated modules were identified by associating module eigengene to case/control status adjusting for age and apoE4 status (*P* < 0.05). KEGG enrichment analysis of significant was conducted using an adjusted *P* value threshold of 0.05. The direction in AD is computed using the case/control coefficient of the model associating module eigengene to case/control status. Modules with significant apoE4:disease interaction effect were identified by adding the interaction term apoE4:disease to the previous model (*P* < 0.05) **B.** Heatmap depicting the degree of module overlap assessed using a hypergeometric test between male and female disease-associated modules. The black module in males had significant overlap (*P* < 0.05) with the pink and yellow modules. Estimate refers to the case/control coefficient in the model module eigengene ∼ age + apoE4 + case/control status. **C.** Hub genes from female disease-associated modules. Hub genes were defined as genes with gene significance (the correlation between the gene expression and case/control status) greater than 0.2 and module membership (the correlation between gene expression and module eigengene) greater than 0.8. Hub genes were restricted to those that were differentially expressed in AD vs control. Protein-protein interactions between hub gene visualization was performed using the STRING v11 database. Edge color represents the type of interaction evidence for protein-protein interaction (cyan: known interaction from curated databases; turquoise: experimentally determined; green: gene-neighborhood predicted interaction; red: gene-fusions predicted interaction; blue: gene co-occurrence predicted interaction; green-yellow: text mining; black: co-expression; light purple: protein homology. **D.** Hub genes among modules with significant apoE4:disease interaction effect. Protein-protein interaction between hub genes was visualized using STRING v11 with edge colors representing the same as in **C**.

Enrichment analysis of disease-associated modules using the 2019 KEGG Human pathway database revealed pathways relevant to AD that were consistent with those identified in the single gene analysis (Figure 3A). For example, in both males and females, an upregulated module was enriched for Akt signaling related pathways and downregulated modules were enriched for oxidative phosphorylation and thermogenesis related pathways, consistent with single gene level analyses.

Notably, several additional pathways not seen through single gene analysis were observed in the network analyses. An upregulated module in both males and females was highly enriched for zinc finger nuclease genes related to Herpes simplex viral infection, consistent with recent work demonstrating Herpes virus infection in AD brains^53^.

Consistent with the single gene analysis, we observed greater number of disease associated modules in females with AD than in males. For example, an upregulated female module was enriched for cell structural processes related to adherens junctions, actin cytoskeleton and axonal guidance. An additional downregulated female module was enriched for neurological signaling pathways including synaptic vesicle exocytosis, aldosterone synthesis and secretion and morphine addiction. Interestingly, an additional female downregulated module was enriched for autophagy and proteolysis pathways, consistent with molecular studies demonstrating decreased autophagy in AD, particularly in females^55^ (Figure 3A).

We also conducted an analysis identifying modules with apoE4:disease interactive effect to understand differential penetrance of the apoE ε4 allele in males and females. In the male gene network, we were unable to identify modules with significant apoE4:disease interactive effect. Interestingly, in the female network, we identified one module that was downregulated (2211 genes) in AD, and two modules (329 genes and 439 genes) that were upregulated in AD and exhibited a significant apoE4:disease interactive effect (Figure 3A). The two upregulated modules (termed pink and purple) were significantly enriched for several zinc finger nuclease genes related to Herpes simplex viral infection. The downregulated module was enriched for metabolic pathways including oxidative phosphorylation and the TCA cycle. Together these results suggest a female-specific network dysregulation involving zinc finger nucleases and metabolic alteration supporting differential apoE4 penetrance in males and females.

There were 102 hub genes among disease associated modules in the female network identified as module membership greater than 0.8, gene significance greater than 0.2, and differentially expressed between AD and controls (Figure 3C; Supplementary Table S7). In contrast, zero hub genes were identified in the male gene network. Protein-protein interaction maps generated by STRING v11 suggest several Ca^+2^- and G protein-dependent interconnected genes including ITPKB, PDGFRB, GNG12, and GNA12 among the female disease associated modules (Figure 3C). Among modules with apoE4:disease interactive effect in females, 35 hub genes were identified, including ITPKB as a highly connected regulator (Figure 3D). For a full list of genes in each module, including hub genes, please refer to Supplementary Table S7).

### Differential Gene Expression in Whole Blood Identifies Stronger Disease Signatures in Females with AD in Comparison to Males

Similar to the brain, we observed distinct AD-associated transcriptomic signatures between males and females with AD in whole blood. We observed a total of 599 differentially expressed genes in females with AD, including 294 upregulated genes and 305 downregulated genes (q < 0.05; Figures 2C-D; Supplementary Table 8). In males, 98 genes were differentially expressed in AD, including 38 upregulated genes and 50 downregulated genes (q < 0.05; Figures 2C-D; Supplementary Table 8). Altogether, 542 genes were uniquely dysregulated in females, including 271 upregulated genes and 271 downregulated genes. In males, 31 genes were uniquely dysregulated, including 15 upregulated genes and 16 downregulated genes. There was a significant overlap of dysregulated genes across males and females with AD (*P* < 0.05; hypergeometric test).

Next, we characterized the transcriptomic signatures observed in the blood of male and female AD patients. Among upregulated genes in female AD patients, we found 14 enriched pathways, many of them relating to components of the innate and adaptive immune system (Table 3; Supplementary Table S9). Several cytokine response elements including STAT5B, STAT6, and IL10RB contributed to enrichment of a number of pathways relating to response to infection (Table 3). Similar to the brain, components of actin cytoskeleton regulation were also dysregulated in females. (Table 3; Supplementary Table S9). Downregulated genes in female AD patients were enriched for a number of metabolism related processes including oxidative phosphorylation and thermogenesis, consistent with the single-gene and network analysis in the brain (Supplementary Table S10).

Similar to the brain analysis, we observed dramatically fewer enriched pathways in males with AD. Among upregulated genes in male AD patients, we did not identify any enriched pathways. Among downregulated genes in male AD patients, components of the proteasome were enriched including PSMD4 and PSMC3 (Table 3; Supplementary Table S11). For a full list of enriched pathways, refer to Supplementary Tables S9-S11.

Lastly, we performed a non-stratified analysis comparing gene expression between AD and control samples irrespective of sex in whole blood. Analyses were adjusted for sex, apoE4 carrier status, age and education. A total of 339 genes were upregulated and 360 genes were downregulated in patients with AD compared to controls (Figure S3B, Supplementary Table S8). Upregulated genes were enriched for several pathways previously implicated in AD, including MAPK signaling, autophagy and NFkB signaling (Supplementary Table S12). In addition, a number of immune related pathways were enriched including tuberculosis, Escherichia coli infection, salmonella infection, and inflammatory bowel disease. Several components of the NFkB cascade and antigen presentation system including NFKBIA, ITGAM, STAT5B, TLR5, TLR4, CD14 and C4A, contributed to this enrichment (Supplementary Table S12). Among downregulated genes, pathways related to protein synthesis and metabolism, including ribosome, proteasome, protein export, thermogenesis, and oxidative phosphorylation were enriched. Included in these pathways were several oxidation phosphorylation related genes including NDUFA9, NDUFA8, COX4I2 (Supplementary Table S13).

### Network Analysis in Whole Blood Identifies a Stronger Disease Signature in Females

We identified five AD-associated modules in females and zero AD-associated modules in males (Figure 4) that met the significance threshold (FDR < 0.05) and were either positively or negatively correlated with case/control status. Among the modules in female samples, three modules including a 483-gene module (termed turquoise), a 129-gene module (termed pink) and 153-gene module (termed black) were upregulated in AD. Two modules including a 270-gene module (termed blue) and 119-gene module (termed magenta) were downregulated in AD (Figure 4A). No modules with significant apoE4:disease interaction effect were found in female or male network analyses from the blood datasets.

**Figure 4:**
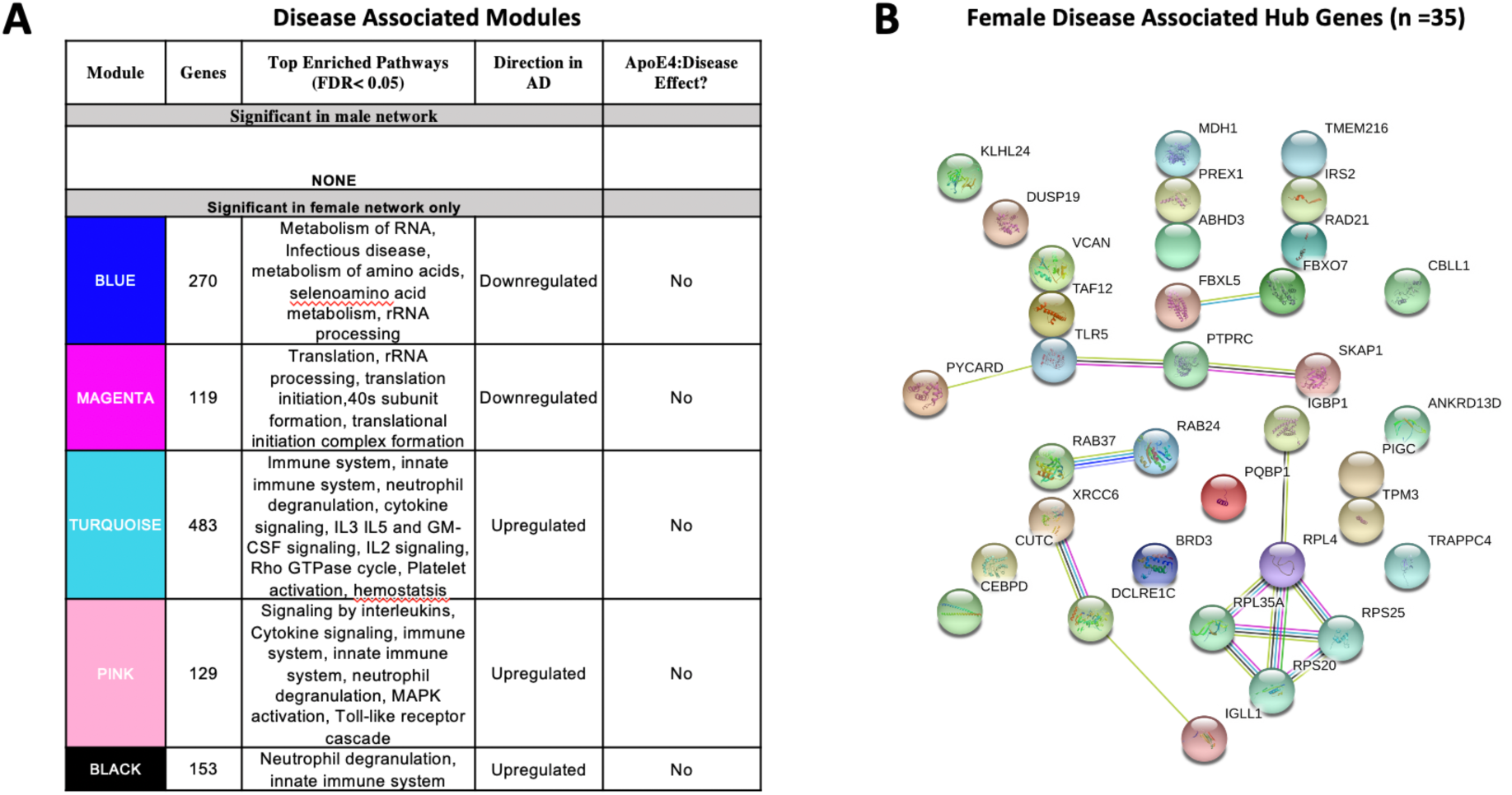
Network Analysis in Whole Blood. WGCNA was used to construct gene network separately for males and females in whole blood. Networks were randomly assigned colors. **A.** A description of the disease-associated gene networks (termed modules) produced using WGCNA. Significant disease-associated modules were identified by associating module eigengene to case/control status adjusting for age and apoE4 status and education (*P* < 0.05). KEGG enrichment analysis of significant was conducted using an adjusted *P* value threshold of 0.05. The direction in AD is computed using the case/control coefficient of the model associating module eigengene to case/control status. Modules with significant apoE4:disease interaction effect were identified by adding the interaction term apoE4:disease to the previous model (*P* < 0.05) **B.** Hub genes from female disease-associated modules. Hub genes were defined as genes with gene significance (the correlation between the gene expression and case/control status) greater than 0.2 and module membership (the correlation between gene expression and module eigengene) greater than 0.8. Hub genes were restricted to those that were differentially expressed in AD vs control. Protein-protein interactions between hub gene visualization was performed using the STRING v11 database. Edge color represents the type of interaction evidence for protein-protein interaction (cyan: known interaction from curated databases; turquoise: experimentally determined; green: gene-neighborhood predicted interaction; red: gene-fusions predicted interaction; blue: gene co-occurrence predicted interaction; green-yellow: text mining; black: co-expression; light purple: protein homology.

Enrichment analysis of disease-associated modules using the 2019 KEGG Human pathway database revealed pathways relevant to AD that were consistent with those identified in the single gene analysis (Figures 4A and 3A). For example, upregulated modules in females were strongly enriched for innate immune system activity, neutrophil degranulation, CSF signaling, IL2 signaling, and cytokine signaling. Consistent with single gene analyses, downregulated modules in females were enriched for metabolic processes including metabolism of RNA and metabolism of amino acids (Figure 4A).

There were 35 hub genes among disease associated modules in the female-specific network identified as module membership greater than 0.8, gene significance greater than 0.2 and differentially expressed between AD and controls (Figure 4B). In contrast, zero hub genes were identified in the male-specific gene network. Protein-protein interaction maps generated by STRING v11 suggest several interconnected genes including the B cell development related protein, IGLL1, and ribosomal proteins RPS20, RPS25, RPL4, and RPL35A (Figure 4B). For a full list of genes in each module, including hub genes, please refer to Supplementary Table S14).

### Comparison of Brain and Blood Transcriptomic Signatures Reveals Common Immune Related Signals in Females

We next identified genes that were commonly dysregulated in both blood and brain (Figure 2E). In females, a total of 23 genes were dysregulated in the brain and blood in the same direction (two downregulated and 21 upregulated). Several genes among the commonly upregulated genes have roles in antigen presentation including TAP1, CTSS, and PTPRC. Enrichment analysis of commonly upregulated genes revealed an enrichment of the KEGG terms primary immunodeficiency, phagosome, and cell adhesion molecules (adjusted *P* < 0.1; Figure 2F). In addition, eight genes were dysregulated but in different directions in the brain and blood including PRKCD, VAMP8, GIMAP7, LAPTM5, HLA-DOA, TNS1, DBI, GIMAP7, TUbA4A (Figure 2E). In contrast, in males we found one upregulated gene, VCAN encoding vesican, dysregulated in both the blood and brain (Figure 2E).

### Cell-type Deconvolution Identifies Sex-specific Immune Cell Dysregulation in Females with AD

Differences in 22 immune blood cell types (Figures 5A-B) were evaluated by deconvolving the transcriptomic signature obtained via meta-analysis of blood studies. Analysis of cell type proportions adjusting for age, sex, and apoE4 status revealed an increase in neutrophils and naïve B cells, and a decrease in M2 macrophages and CD8+ T cells in AD patients compared to controls in pooled male and female samples (Figure 5C, FDR *P* <0.05). Among females with AD, relative to controls, we observed an increase in neutrophils and naïve B cells and a decrease in M2 macrophages, memory B cells, and CD8+ T cells in AD samples (Figure 5C, FDR *P* <0.05). Interestingly, among males with AD, we did not find any significant differences in immune cell proportions compared to controls.

**Figure 5:**
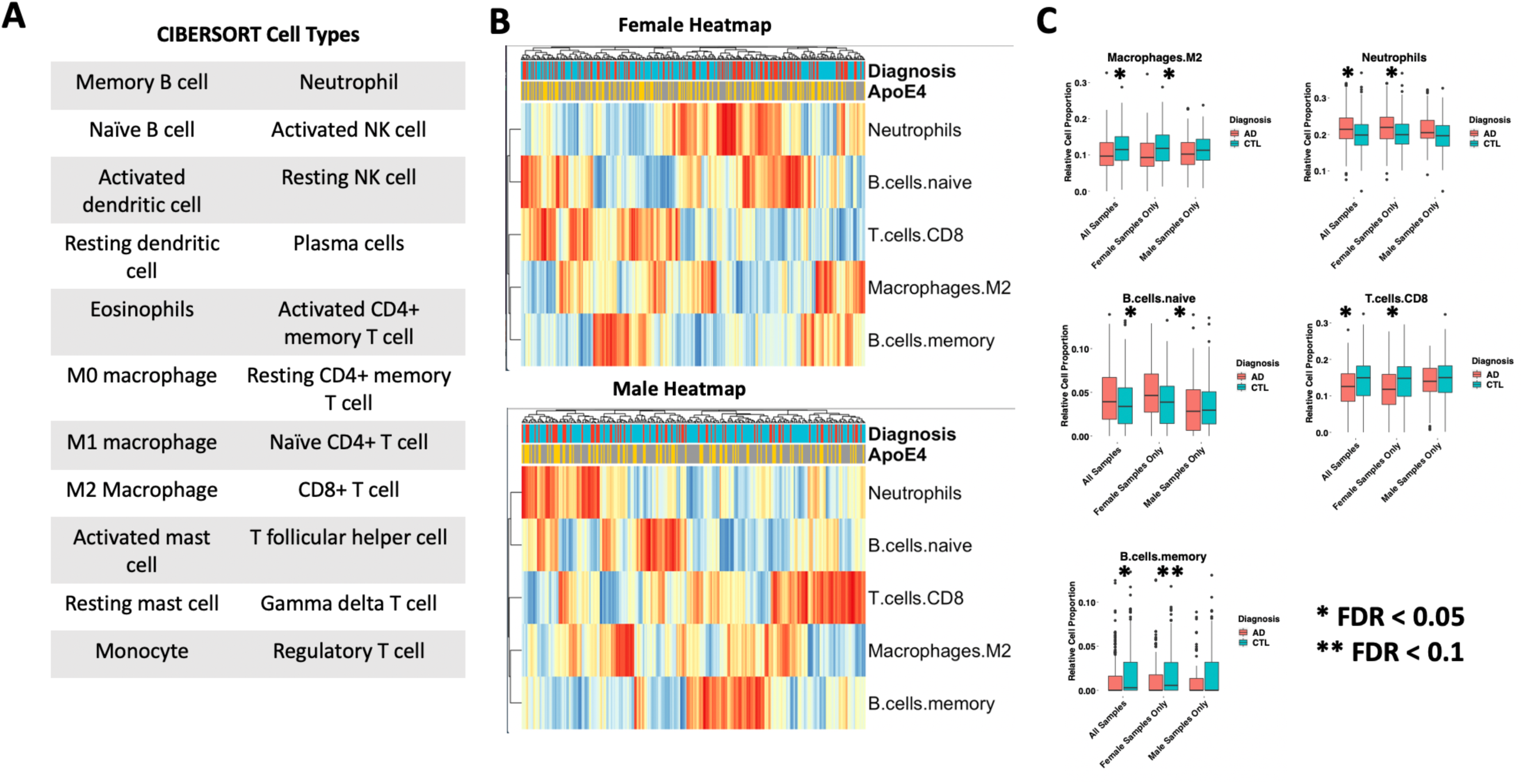
Cell Type Analysis in Whole Blood. **A.** Cell types included in the panel of 22 reference cell types in CIBERSORT. **B.** Heatmap depicting cell type expression between cases and controls. apoE4 carrier status, sex, and case/control status is annotated for each sample. Only cell types that are significantly different between cases and controls in pooled male and female, male-only or female-only analyses are shown. **C.** Bar charts depicting cell type expression for individual cell types that are significantly between cases and controls in pooled male and female, male-only or female-only analyses are shown. Significance was assessed by associated cell type proportion to case/control status, adjusting for age, sex (in the pooled male and female model) and apoE4 status. *P* < 0.05 was deemed significant.

### Sex-specific Transcriptomic Data Improves AD Classification Accuracy

To assess the value of sex-specific transcriptomic data in developing a blood-based classifier in AD, we trained a linear SVM model to classify AD patients controls using the transcriptomic signature obtained via meta-analysis of blood studies. We trained a ‘clinical model’ with age, sex, education, and apoE4 status and a ‘clinical + molecular model’ with age, sex, education, apoE4 status, and blood transcriptomic data. Using pooled male and female samples, the ‘clinical + molecular model’ achieved a higher AUROC compared to the ‘clinical model’ (AUROC = 0.88 for ‘clinical + molecular model’; AUROC = 0.77 for ‘clinical model’) on a test set composed of 25% of samples (Figures 6A and S4A).

**Figure 6:**
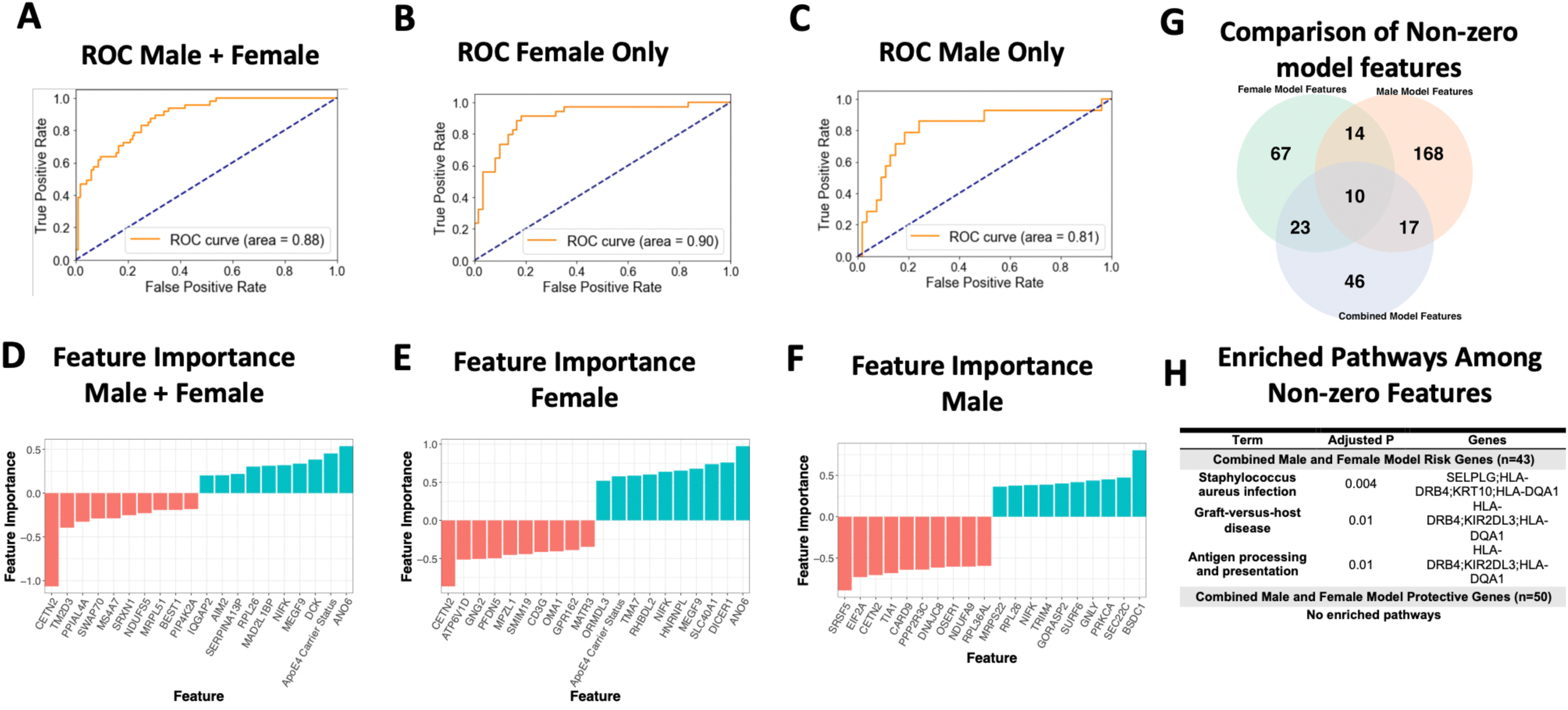
Linear SVM Clinical + Molecular Model in Whole Blood. **A-C.** Receiver operating characteristic (ROC) curves depicting performance of each linear SVM model on a test set composed of 25% of samples. Features include gene expression data obtained via meta-analysis, age, sex, education, and apoE4 status. Three models were fit for male and female pooled samples (**A)**, female samples only (**B**), and male samples only (**C**). **D-F.** Feature importance plots for features with non-zero importance in the combined male and female model (**D)**, female model (**E**), and male model (**F**). A positive feature importance means that the expression of that feature increases the likelihood of being classified as AD (risk factor). A negative feature importance means that expression of the feature expression reduces the likelihood of being classified as AD (protective factor). **G.** Comparison of non-zero features between combined male and female model, female model and male model.

Interestingly, a model trained with only female data achieved a higher AUROC (‘clinical + molecular model’: 0.90 and ‘clinical model’: 0.86; Figures 6B and S4B) than the pooled male and female model. In contrast, a model trained with only male data obtained a lower AUROC (‘clinical + molecular’ model 0.81 and ‘clinical model’ 0.83; Figures 6C and S4C) than the pooled male and female model.

Figures 6G-H summarizes shared features between models. In all simple models (pooled male and female, female only, and male only), age and apoE4 status had a positive feature importance while education had a negative feature importance. A positive feature importance means that the expression of that feature increases the likelihood of being classified as AD (termed risk factor). A negative feature importance means that expression of the feature expression reduces the likelihood of being classified as AD (termed protective factor). In the female ‘clinical + molecular model’, 57 features, including known risk factors including apoE4 and age, had a positive feature importance (Supplementary Table S15). In addition, 50 features had negative feature importance. Among these were education and previously implicated AD risk genes including CETN2 (Supplementary Table S15). In the male ‘clinical + molecular model’, 103 features, including apoE4, had positive feature importance. (Supplementary Table S16). In addition, 105 features, including education, had negative feature importance (Supplementary Table S16).

Altogether, we observed a significant overlap (*P* < 0.001, hypergeometric test) in features with non-zero feature importance between the pooled male and female ‘clinical + molecular model’ and female ‘clinical + molecular model’; female ‘clinical + molecular model’ and male ‘clinical + molecular model’; and pooled male and female ‘clinical + molecular model’ and male ‘clinical + molecular model’ (Figure 6G).

Functional annotation of features with a non-zero feature importance was performed via enrichment analysis using the 2019 KEGG database of human pathways. Among features with non-zero feature importance, we did not identify any enriched biological pathways in the male only and female only complex models. In the male and female pooled complex model, features with positive feature importance (risk factors), were enriched for staphylococcus aureus infection, graft-vs-host disease, and antigen presentation and processing KEGG pathways (adjusted *P* < 0.05; Figure 6H). The HLA genes HLA-DRB4 and HLA-DQA1 contributed to this enrichment. In addition, the P-selection glycoprotein ligand-1 gene (SELPLG) and killer cell immunoglobulin-like receptor (KIR2DL3) also contributed to enrichment, suggesting a role for leukocyte recruitment and natural killer cell activity in AD pathology.

## DISCUSSION

In this study, through computational analysis of publicly available gene expression datasets from brain and blood samples, we evaluated AD at the transcriptome level using single gene and network approaches to gain insight into the mechanisms underlying sex and apoE4-genotype based differences in AD. We also evaluated how including sex-specific transcriptomic data from blood samples with clinical data would affect the performance of a machine learning classifier for AD diagnostics.

Our characterization of brain transcriptomic signatures revealed, among upregulated genes in the brains of both females and males with AD, an enrichment of pathways related to components of the innate and adaptive immune systems as well as the MAPK signaling pathway. This result is consistent with past findings where the brain’s immune system has been indicated as a major component of AD pathogenesis^60, 61^. Additionally, MAPKs, enzymes that play critical roles in cellular signaling, have also been implicated as accelerators of AD development^62^. Overall, findings from our brain transcriptome analysis provide supporting evidence for therapeutics currently being explored for AD, such as p38 MAPK inhibitors^67^, and suggest that possible treatments targeting the MAPK pathway may have a greater effect in females with AD.

Interestingly, from our differential expression analysis, we found a nearly two-fold greater total number of dysregulated genes in the brain transcriptome that met our significance cutoff for females with AD compared to males with AD (974 vs 509, respectively). Many of these genes are in pathways related to antigen presentation and processing, complement activation, suggesting a female-specific role of neuroinflammation in the pathogenesis of AD. Additionally, for downregulated genes in AD patients, we observed enrichment of neurological signaling pathways in females only and no enriched pathways in males.

Through network analysis, we identified more AD-associated modules in the brain transcriptome of females than males. Enrichment analysis of AD-associated modules also revealed some pathways that were enriched in both sexes, including an upregulated module for a PI3/Akt signaling related pathway and downregulated modules for oxidative phosphorylation and thermogenesis related pathways. Unique to females, we observed upregulated modules associated with cell structural processes (adherens junctions, actin cytoskeleton and axonal guidance) and HSV infection-related zinc finger nuclease genes, as well as a downregulated module for neurological signaling pathways, autophagy and proteolysis.

Upon performing hub gene analysis, we identified hub genes in female disease-associated modules but were unable to identify male disease associated hub genes. These female hub genes consisted of several potentially interconnected genes including ITPKB, PDGFRB, GNG12, and GNA12. In our subsequent analysis to assess an apoE4:disease interaction effect, we identified three modules, one of which was significantly enriched for HSV infection-related zinc finger nuclease genes as well as containing the ITPKB hub gene as a highly connected regulator. These results suggest zinc finger nucleases as a potential mechanism underlying sex-associated differential penetrance of apoE4 in AD.

Our findings suggest a neuroinflammatory model of AD pathogenesis in females with dysregulation in components of the adaptive and innate immune system including antigen presentation and processing and complement activation and genes including MAPK and ITPKB. It has been postulated that accumulation of damage from HSV infection and major neuroinflammatory effects can lead to the development of AD, and that apoE4 carriers suffer either greater viral damage or have poorer repair of such damage^54^. Previous studies have demonstrated that ITPKB expression is increased in human AD brains and exacerbates AD pathology in an animal model^64^. Our brain transcriptome findings for females with AD, including downregulation of autophagy and proteolysis pathways, upregulation of pathways related to the immune system and HSV infection, as well as ITPKB as a hub gene, particularly in female apoE4 carriers, highlight specific gene-encoded processes in the brain that may be more involved in AD for women than for men.

Similar to our brain findings, in analysis of blood transcriptomes, we observed more dysregulated genes in the blood of females with AD than in males with AD. Further characterization of these transcriptomic signatures revealed, among upregulated genes, enrichment in only females with AD of pathways related to components of the innate and adaptive immune systems as well as actin cytoskeleton regulation; however, for downregulated AD genes, we observed enriched metabolic pathways (oxidative phosphorylation and thermogenesis) in females and enriched pathways for protein homeostasis in males.

Through network analysis, we identified AD-associated modules and hub genes in the female blood transcriptome but not in males. In the blood of females with AD, upregulated modules were strongly enriched for innate immune system activity (neutrophil degranulation, CSF signaling, IL2 signaling, and cytokine signaling). Consistent with single gene analyses, female downregulated modules were enriched for metabolic processes (e.g. metabolism of RNA and amino acids). Hub genes identified in the blood of females with AD include those related to immunity (the B cell development related protein, IGLL1) and viral RNA translation (ribosomal proteins RPS20, RPS25, RPL4, and RPL35A).

In addition to neuroinflammation’s role in AD, dysregulation of the immune system outside of the brain has also been noted to be a factor in AD^65^. Our findings feature specific gene-encoded processes in peripheral blood cells that may be more involved in AD for women than for men. Furthermore, our cell-type deconvolution analysis revealed dysregulation of peripheral immune cells uniquely in females with AD and not males with AD.

When including blood transcriptomic features with clinical features (age, sex, education, and apoE4 status) to train a machine learning prediction model of AD, our model performed better with these additional molecular features than without (AUROC: 0.88 vs 0.77, respectively). The performance of this model also improved when trained with only female data (clinical + molecular model AUROC: 0.90 and clinical model AUROC: 0.86) and worsened when trained with only male data (molecular model AUROC: 0.81 and clinical model AUROC: 0.83) than with pooled male and female model. This finding suggests that the molecular changes in females compared to males are better able to model AD-related changes. Further, given the distinct transcriptomic signature observed in males and females, stratifying by sex may aid future efforts to identify biomarkers in AD.

Diagnostic tests currently available for AD, including Aβ position emission tomography (PET), lack accuracy or are implemented through invasive and painful procedures such as lumbar puncture^56–59^. Diagnostic tests for AD that are more accurate and less invasive are worthwhile for preventing undue uncertainty and physical discomfort experienced by patients. Our machine learning AD prediction model based on clinical and blood transcriptomic features has the potential to complement currently available clinical AD diagnostic tests, and improve the accuracy of these tests, particularly for women, with minimal additional discomfort for patients.

Based on the nature of our analyses, there are a number of limitations to note. We analyzed publicly available datasets, which were limited in sample size and contained annotation differences. This provided challenges in selecting cases from controls and restricted our ability to answer certain questions. For instance, the Allen Brain Atlas dataset provided only a binary classification for apoE (apoE4: Y/N). This confined our analysis to only look at the presence of apoE4, instead of looking at difference across different genotype combinations. Next, we did not stratify our analysis by age or disease stage, so we cannot describe whether these transcriptomic signatures differ with age or disease severity. Additionally, since we aggregated bulk tissue from different brain regions in our analysis, we cannot infer sex differences across brain region. Consequently, using bulk tissue transcriptomics reduces our resolution of the more complex interactions and contributions of different brain cell types in AD. Future approaches to better characterize sex-specific changes in AD would involve stratification by brain regions, age and disease stage, apoE genotype, as well as an analysis of single cell AD datasets.

In conclusion, the major finding of this study is a distinct, sex-specific transcriptomic signature in the brains and whole blood of patients with AD. Gene expression meta-analysis and network-based analyses revealed an immune signature in the brains and whole blood of females with AD that was absent in males. Our analyses also revealed more pronounced neurosignaling and metabolism signatures in the brains whole blood of females with AD than in males with AD. Stratification by sex improved machine-learned based classification of AD using whole-blood transcriptomic data. Results from this work will help to better understand molecular etiologies underlying sex differences in AD and pave the way for sex-specific biomarker and therapeutic development in AD.

## Acknowledgements

This work was in part supported by NIA 1R0AG057683 and R01AG060393. Data collection and sharing for this project was funded by the Alzheimer’s Disease Neuroimaging Initiative (ADNI) (National Institutes of Health Grant U01 AG024904) and DOD ADNI (Department of Defense award number W81XWH-12-2-0012). ADNI is funded by the National Institute on Aging, the National Institute of Biomedical Imaging and Bioengineering, and through generous contributions from the following: AbbVie, Alzheimer’s Association; Alzheimer’s Drug Discovery Foundation; Araclon Biotech; BioClinica, Inc.; Biogen; Bristol-Myers Squibb Company; CereSpir, Inc.; Cogstate; Eisai Inc.; Elan Pharmaceuticals, Inc.; Eli Lilly and Company; EuroImmun; F. Hoffmann-La Roche Ltd and its affiliated company Genentech, Inc.; Fujirebio; GE Healthcare; IXICO Ltd.; Janssen Alzheimer Immunotherapy Research & Development, LLC.; Johnson & Johnson Pharmaceutical Research & Development LLC.; Lumosity; Lundbeck; Merck & Co., Inc.; Meso Scale Diagnostics, LLC.; NeuroRx Research; Neurotrack Technologies; Novartis Pharmaceuticals Corporation; Pfizer Inc.; Piramal Imaging; Servier; Takeda Pharmaceutical Company; and Transition Therapeutics. The Canadian Institutes of Health Research is providing funds to support ADNI clinical sites in Canada. Private sector contributions are facilitated by the Foundation for the National Institutes of Health (www.fnih.org). The grantee organization is the Northern California Institute for Research and Education, and the study is coordinated by the Alzheimer’s Therapeutic Research Institute at the University of Southern California. ADNI data are disseminated by the Laboratory for Neuro Imaging at the University of Southern California.

## Author Contributions

MDP, MS conceived and designed the study. MDP, AT, and BG extracted data from publicly available data sources used in the study. MDP, JKW, IP, AG completed all analyses and produced figures. MDP, MS, TO, and SB wrote the manuscript. MDP, MS, TO, SB, AT, KZ, BG, and YH critically revised the manuscript for important intellectual content. All authors commented and approved the manuscript.

## Competing Interests

Y.H. is a co-founder and scientific advisory board member of E-Scape Bio, Inc. and GABAeron, Inc. M.S. is on the scientific advisory board for twoXAR. The other authors declare no competing interests.

## Supplementary Figure Legends

**Figure S1: Brain Data PCA**

Principal component plots (PCA) of brain samples before (top) and after batch (bottom) correction. PCA plots depict principal component 1 and 2 and are colored by study (left), sex (middle) and apoE4 status (right).

**Figure S2: Blood Data PCA**

Principal component plots (PCA) of blood samples before (top) and after batch (bottom) correction. PCA plots depict principal component 1 and 2 and are colored by study (left), sex (middle) and apoE4 status (right).

**Figure S3: Non-stratified Differential Gene Expression**

**A.** Volcano plot depicting fold changes and p values from samples in the brain. Analyses were adjusted for sex, apoE4 status, and age. An adjusted *P* value < 0.05 and log FC > 1.2 was deemed significant. In the brain, a total of 662 genes were upregulated and 430 genes were downregulated in patients with AD compared to controls. **B.** Volcano plot depicting fold changes and *P* values from samples in the blood. Analyses were adjusted for sex, apoE4 status, age, and education. An adjusted *P* value < 0.05 was deemed significant. In blood, 339 genes were upregulated and 360 genes were downregulated in patients with AD compared to controls.

**Figure S4: Linear SVM Clinical Model**

**A-C.** Receiver operating characteristic (ROC) curves depicting performance of each linear SVM model on a test set composed of 25% of samples. Features include age, sex, education, and apoE4 status. Three models were fit for male and female pooled samples (**A)**, female samples only (**B**), and male samples only (**C**). **D-F.** Feature importance plots for features with non-zero importance in the combined male and female model (**D)**, female model (**E**), and male model (**F**). A positive feature importance means that the expression of that feature increases the likelihood of being classified as AD (risk factor). A negative feature importance means that expression of the feature expression reduces the likelihood of being classified as AD (protective factor).

