## Supplemental Figures for "Sex-Specific Cross Tissue Meta-Analysis Identifies Immune Dysregulation in Women with Alzheimer’s Disease"

**Table S1: Differential expression fold changes and p-values in the brain**

**Table S2: Pathway Enrichment Among Upregulated Genes in Female Analysis in the Brain**

**Table S3: Pathway Enrichment Among Downregulated Genes in Female Analysis in the Brain**

**Table S4: Pathway Enrichment Among Upregulated Genes in Male Analysis in the Brain**

**Table S5: Pathway Enrichment Among Upregulated Genes in Combined Male and Female Analysis in the Brain**

**Table S6: Pathway Enrichment Among Downregulated Genes in Combined Male and Female Analysis in the Brain**

**Table S7: Module Membership in Females in Brain**

**Table S8: Differential expression fold changes and p-values in whole blood**

**Table S9: Pathway Enrichment Among Upregulated Genes in Female Analysis in Whole Blood**

**Table S10: Pathway Enrichment Among Downregulated Genes in Female Analysis in Whole Blood**

**Table S11: Pathway Enrichment Among Downregulated Genes in Male Analysis in Whole Blood**

**Table S12: Pathway Enrichment Among Downregulated Genes in Combined Male and Female Analysis in Whole Blood**

**Table S13: Pathway Enrichment Among Upregulated Genes in Combined Male and Female Analysis in Whole Blood**

**Table S15: Module Membership in Males in Whole Blood**

**Table S16 Linear SVM Feature Importance for Clinical + Molecular Model in Females**

**Table S17 Linear SVM Feature Importance for Clinical + Molecular Model in Males**

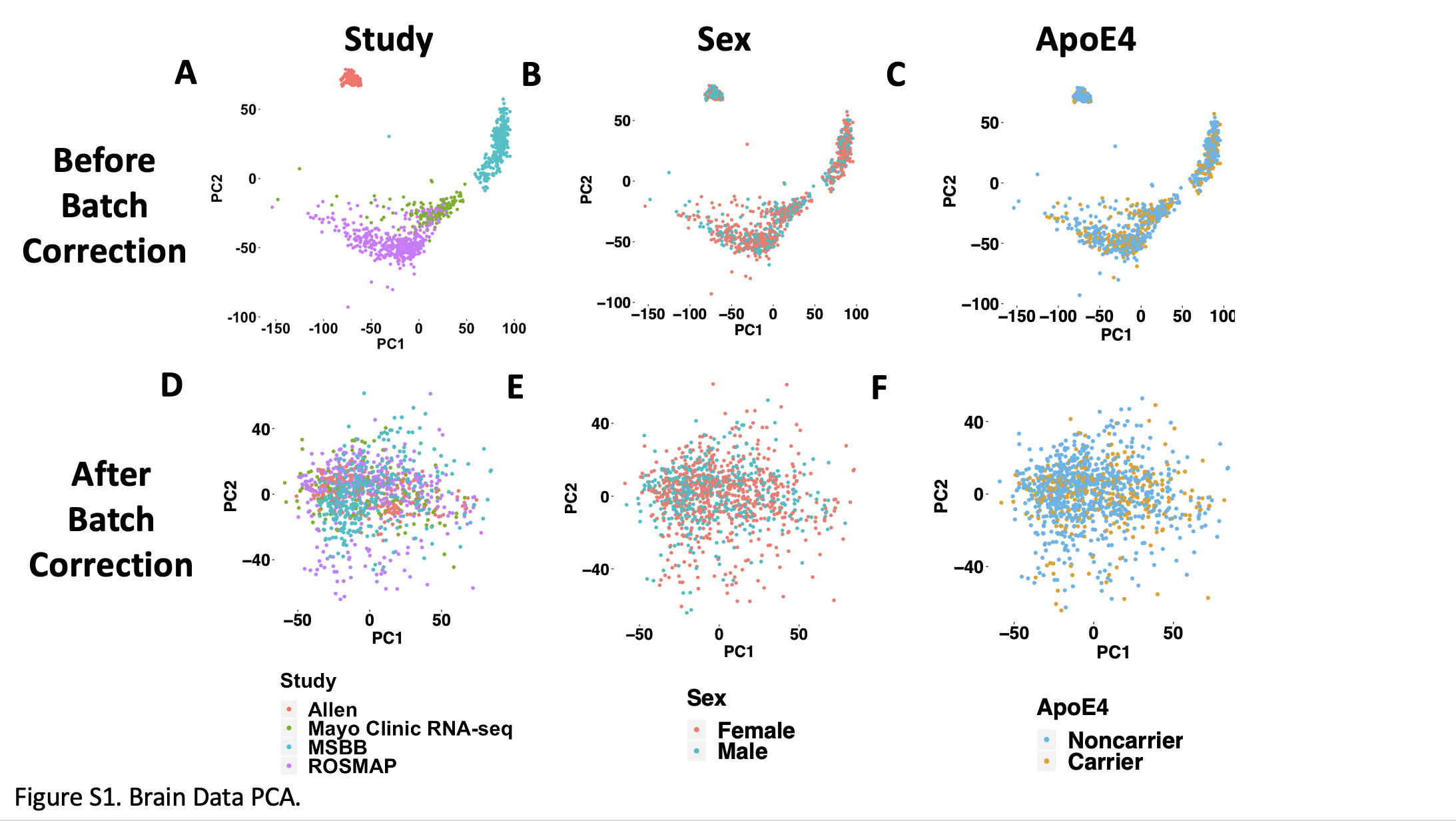

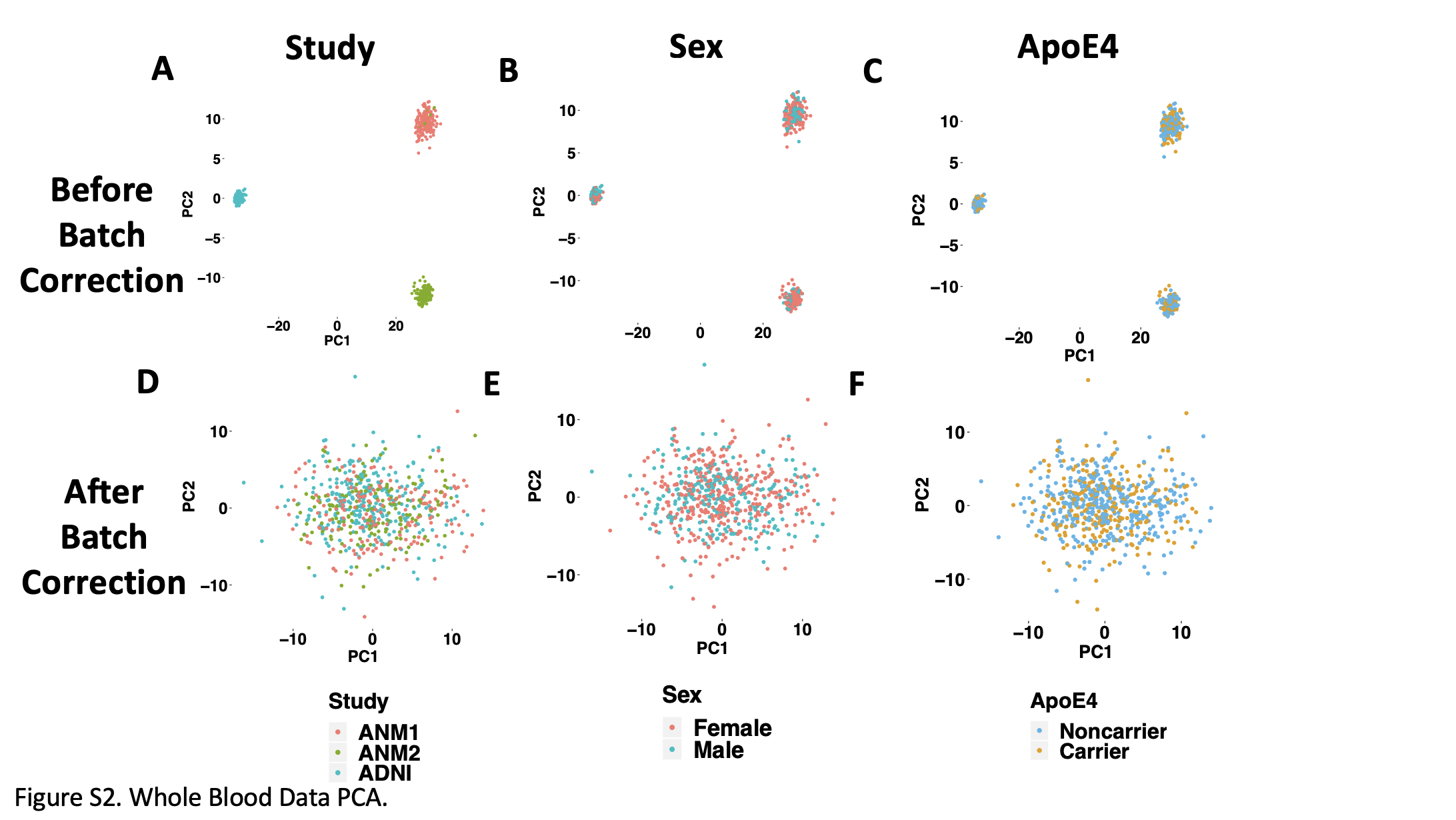

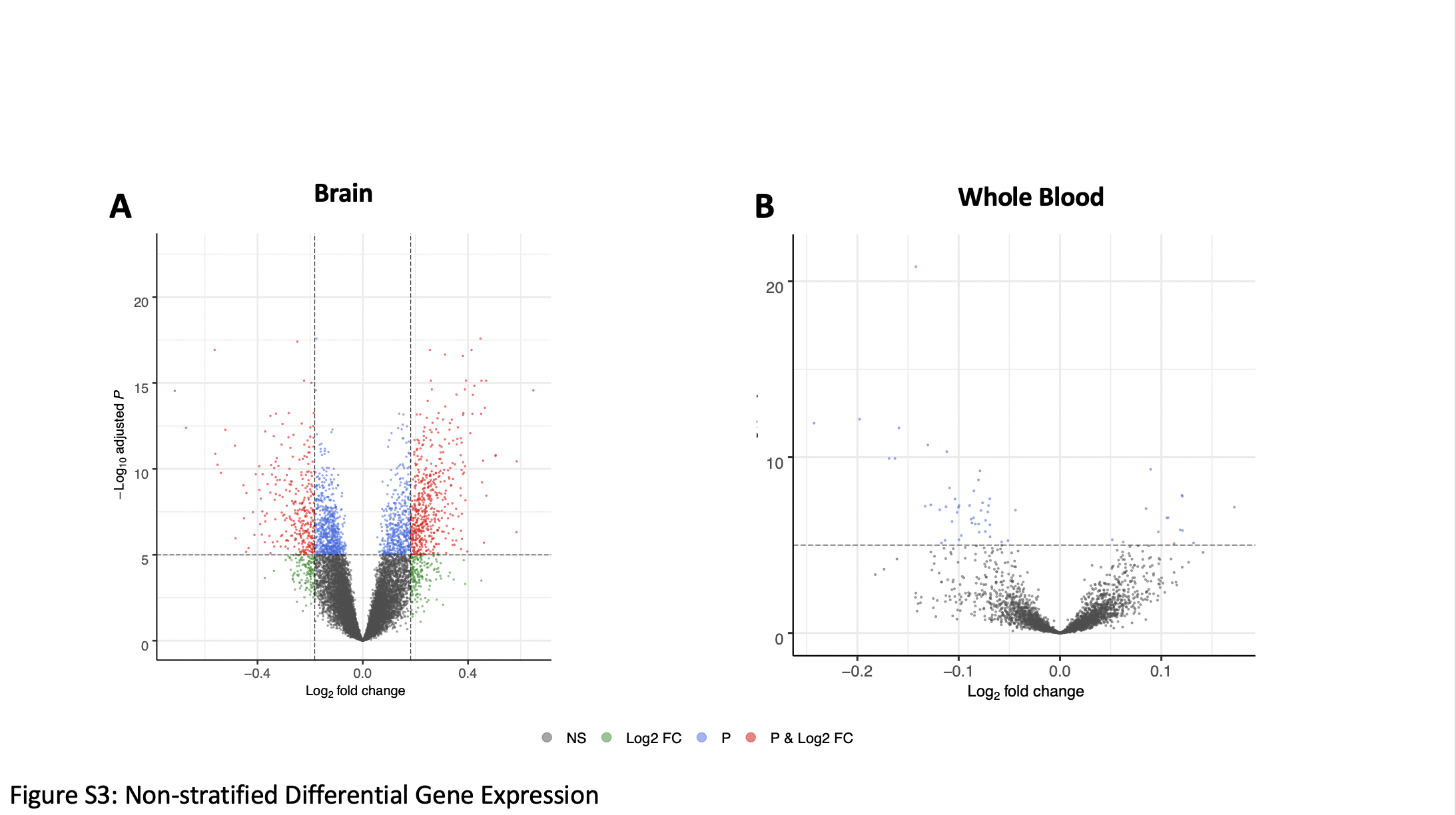

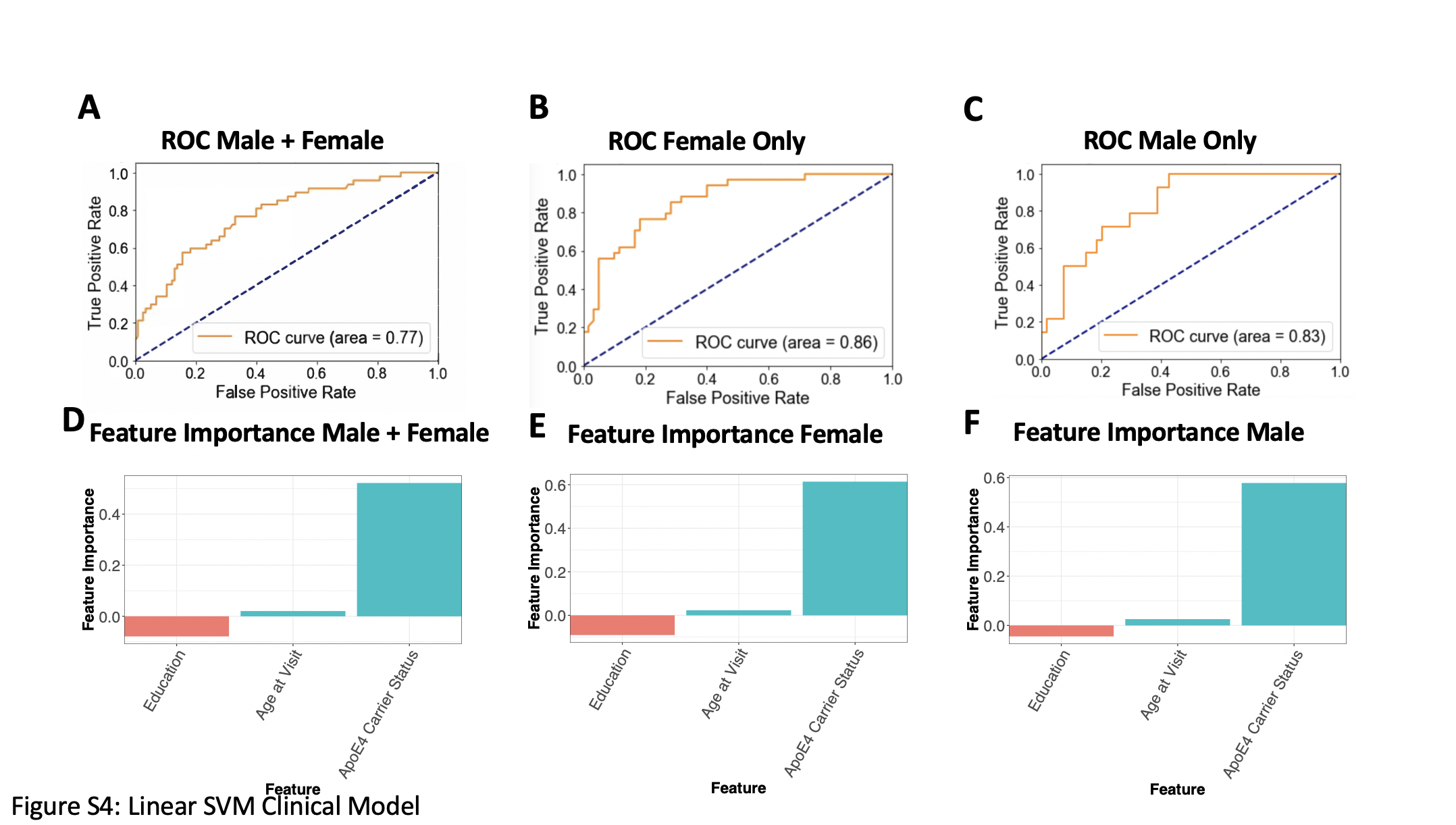
